# CopulaNet: Learning residue co-evolution directly from multiple sequence alignment for protein structure prediction

**DOI:** 10.1101/2020.10.06.327585

**Authors:** Fusong Ju, Jianwei Zhu, Bin Shao, Lupeng Kong, Tie-Yan Liu, Wei-Mou Zheng, Dongbo Bu

**Affiliations:** Key Lab of Intelligent Information Processing, Big-data Academy, Institute of Computing Technology, Chinese Academy of Sciences, Beijing, China; University of Chinese Academy of Sciences, Beijing, China; Microsoft Research Asia, Beijing, China; Institute of Theoretical Physics, Chinese Academy of Sciences, Beijing, China

## Abstract

Protein functions are largely determined by the final details of their tertiary structures, and the structures could be accurately reconstructed based on inter-residue distances. Residue co-evolution has become the primary principle for estimating inter-residue distances since the residues in close spatial proximity tend to co-evolve. The widely-used approaches infer residue co-evolution using an indirect strategy, i.e., they first extract from the multiple sequence alignment (MSA) of query protein some handcrafted features, say, co-variance matrix, and then infer residue co-evolution using these features rather than the raw information carried by MSA. This indirect strategy always leads to considerable information loss and inaccurate estimation of inter-residue distances. Here, we report a deep neural network framework (called CopulaNet) to learn residue co-evolution directly from MSA without any handcrafted features. The CopulaNet consists of two key elements: *i*) an encoder to model context-specific mutation for each residue, and *ii*) an aggregator to model correlations among residues and thereafter infer residue co-evolutions. Using the CASP13 (the 13th Critical Assessment of Protein Structure Prediction) target proteins as representatives, we demonstrated the successful application of CopulaNet for estimating inter-residue distances and further predicting protein tertiary structure with improved accuracy and efficiency. Head-to-head comparison suggested that for 24 out of the 31 free modeling CASP13 domains, ProFOLD outperformed AlphaFold, one of the state-of-the-art prediction approaches.

---

Proteins play critical roles in a wide-range of biological processes including catalyzing metabolic reactions, responding to stimuli, and transporting molecules, and these biological activities are largely determined by protein tertiary structures [1]. Protein tertiary structures can be experimentally determined using nuclear magnetic resonance, X-ray crystallography, and cryogenic electron microscopy [2]; however, these technologies are usually costly and time-consuming and cannot keep pace with the rapid accumulation of protein sequences [3]. Thus, the computational approaches for predicting protein structures purely from residue sequences are highly desirable [4, 5].

Major progresses have been made during previous years in protein structure prediction and inter-residue contacts/distances have played important roles. Most of the recent protein structure prediction approaches, such as AlphaFold [6] and trRosetta [7], employ roughly the same three-step diagram: *i*) estimating inter-residue contacts/distances; *ii*) constructing an energy function based on the estimated contacts/distances; and *iii*) building the tertiary structure such that the energy is minimized. This diagram has been shown to be successful when the estimated inter-residue contacts/distances are sufficiently accurate.

The state-of-the-art approaches to estimating the inter-residue contacts/distances share the same cornerstones: constructing multiple sequence alignment (MSA) for target protein and then exploiting MSA using the principle of residue co-evolution [8–10]. The rational underlying this principle is that two residues in close spatial proximity always tend to co-evolve; thus, in turn, the co-evolution could be exploited to accurately estimate contacts/distances between residues. Here, the co-evolution relationship is inferred from residue correlations shown in MSA.This principle, however, is always hindered by the indirect correlations among residues: the indirect correlations could lead to transitivity in residue spatial proximity and thereafter incorrect estimation of inter-residue distances. To extract the direct correlations, a variety of direct coupling analysis (DCA) methods have been proposed using precision matrix (the inverse of covariance matrix) or Potts model [11–15]. Currently the DCA technique is widely used for estimating inter-residue contacts, especially combined with deep neural networks for further refinement. For example, both AlphaFold [6] and RaptorX [16] rely on the inter-residue contacts predicted by CCMpred, a DCA-based approach using the Potts model [17].

Although the DCA technique has been shown to be effective in estimating inter-residue distances, it still suffers from several drawbacks. An outstanding drawback is the considerable information loss after transforming MSAs into handcrafted features, say covariance matrices. In fact, the DCA technique is founded on the premise that the direct correlations between two residues can be modeled using pair-wise statistics such as covariance [9, 10]. However, this premise does not always hold. We demonstrated this possibility using two artifactual proteins *P*_1_ and *P*_2_ as counterexamples (Figure 1). In protein *P*_1_, two residues *R*_1_ and *R*_2_ are close, whereas in protein *P*_2_, they are far from each other. Despite the substantial difference in the constructed MSAs for *P*_1_ and *P*_2_, the covariance matrices calculated from these MSAs are completely identical, causing the DCA technique to give identical distance estimations for proteins *P*_1_ and *P*_2_. In fact, for these two MSAs, any statistic of a single residue, or pairwise statistics of two residues, cannot distinguish them. Thus, a more effective way to extract direct correlation from MSAs is highly desirable.

**Figure 1.**
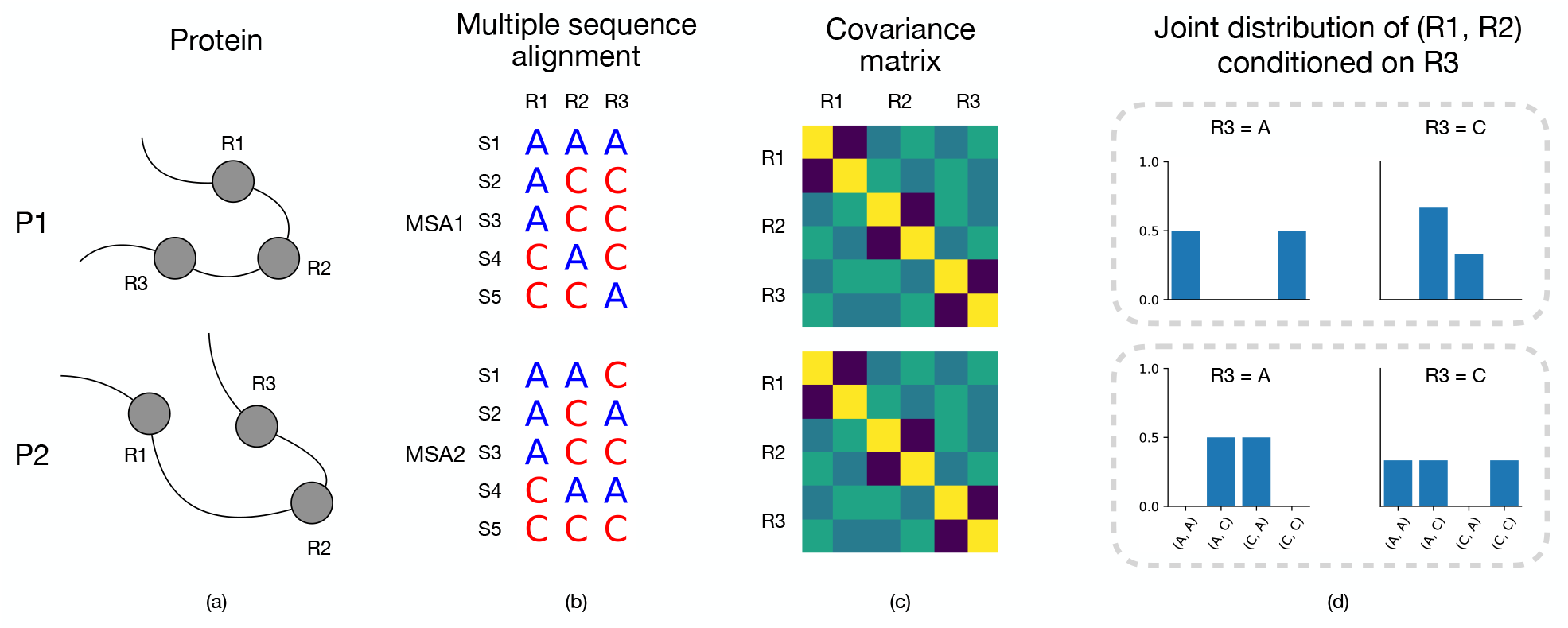
The limitation of the covariance-based methods in estimating inter-residue distances. (a) Two artifactual proteins *P*_1_ and *P*_2_. In protein *P*_1_, two residues *R*_1_ and *R*_2_ are close, whereas in protein *P*_2_, they are far from each other. (b) The MSAs constructed for the two proteins show considerable difference. (c) The covariance matrices calculated from these two MSAs are totally identical; thus, the covariance-based methods give the same estimation of inter-residue distances for protein *P*_1_ and *P*_2_. This is contradict to the real inter-residue distances. (d) Unlike the covariance matrices, the conditional joint-residue distribution *P* (*R*_1_, *R*_2_|*R*_3_) could effectively distinguish these two MSAs

Unlike covariance matrices, the conditional joint-residue distribution could effectively describe the direct correlation between residues (Fig. 1d). Here, we report an approach (called ProFOLD) for inter-residue distance estimation through learning the conditional joint-residue distributions directly from MSAs rather than the handcrafted features such as covariance matrices.

The core of our approach is a novel deep neural network framework CopulaNet, which consists of three key elements: an MSA encoder, a coevolution aggregator, and a distance estimator. The MSA encoder processes each homologue protein in MSA individually, and embeds each residue to represent its context-specific mutations observed from the homologue protein. For any two residues, the aggregator first calculates the outer product of their embeddings for each homologue protein, then aggregates the out products acquired from all homologue proteins using average pooling, and finally yields a measure of the coevolution between the two residues. Based on the obtained residue coevolution, we use a two-dimensional residual network to estimate distance for any residue pairs. Further, ProFOLD transforms the estimated distances into a potential function, and searches the tertiary structural conformation with the minimal potential.

We demonstrated the concept of ProFOLD using protein T0992-D1 as an example, applied it to predict structures for the CASP13 target proteins as representatives, and compared it with the state-of-the-art prediction approaches.

## Results and discussions

### Approach summary

The ProFOLD approach is summarized in Figure 2. Using a CASP13 target protein T0992-D1 as an example, we demonstrate the concept and main steps of ProFOLD for protein structure prediction.

**Figure 2.**
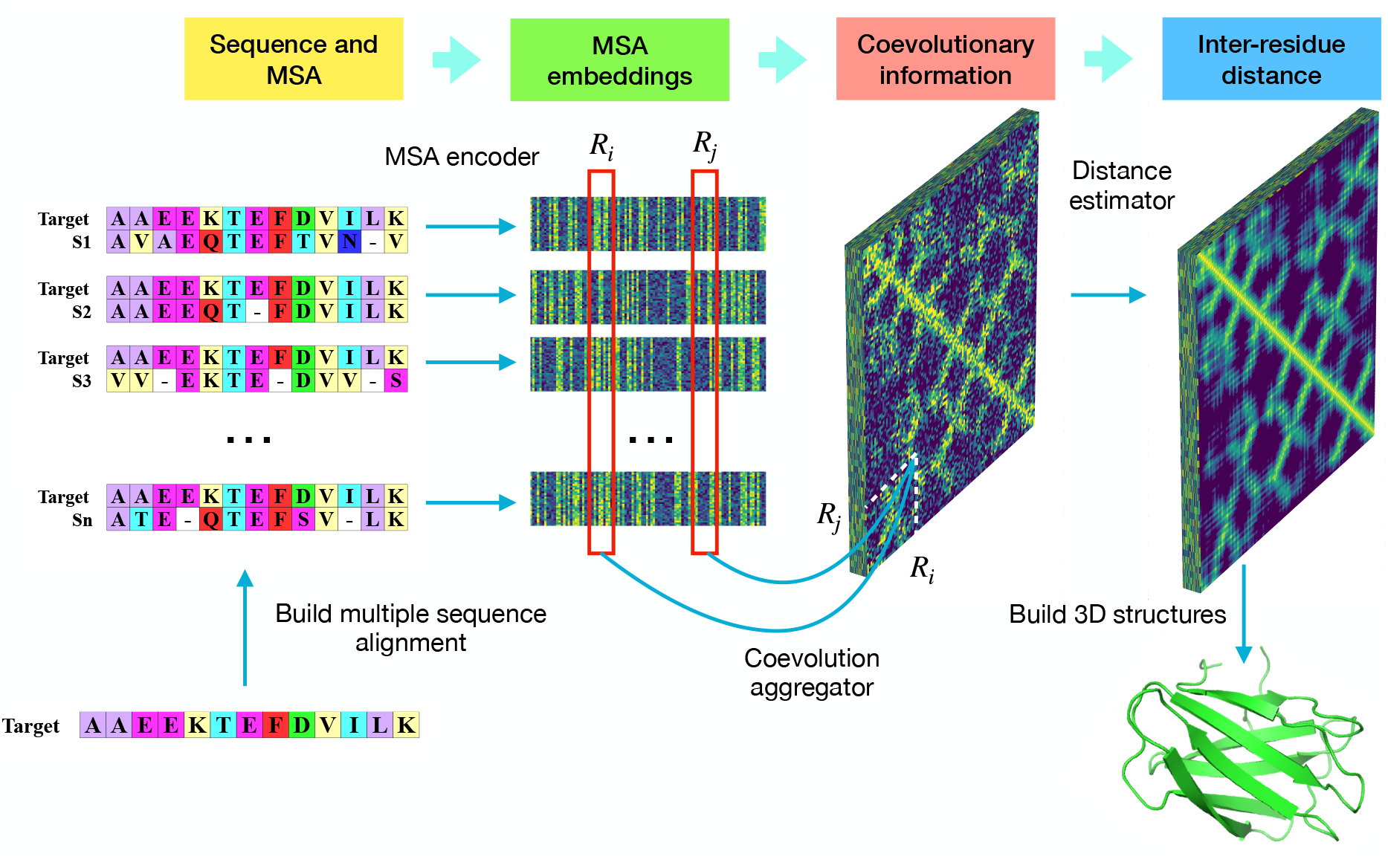
Predicting protein tertiary structure using ProFOLD. Here, we use protein T0992-D1 as an example to describe the main steps of ProFOLD. Only the first 13 residues are shown here for the sake of clear description. First, we search homologue proteins (2807 in total) for this protein and construct MSA accordingly. Then we apply CopulaNet to infer residue co-evolution directly from the MSA. CopulaNet uses an MSA encoder to model the mutation information for each residue of the target protein, and then uses a co-evolution aggregator to measure the residue co-mutations. According to the acquired residue co-evolutions, the distance estimator estimates inter-residue distances. Finally, we transform the estimated distance distributions into a potential function, and then search for the structure conformation with the minimal potential. ProFOLD reports the structural conformation with sufficiently low potential as the final prediction result (TMscore: 0.84)

Protein T0992-D1 consists of a total of 107 residues (only the first 13 residues are shown here for the sake of clear description). For this protein, we first identify its homologues through searching it against protein sequence databases including uniclust30, uniref90 and metaclust50. The obtained homologue proteins (2807 in total) are organized into an MSA. Next, we apply CopulaNet to infer inter-residue distances directly from the constructed MSA. Here, we infer the distribution of inter-residue distance over pre-defined 37 bins, rather than a single distance value. Four examples of these distributions are shown in Supplementary Figure 1. In the case of residues LEU32 and TYR70, the most likely distance interval was predicted to be (7.5Å, 8Å), which covers the true distance 7.83Å. Finally, we transform the estimated distance distributions into a potential function, and then search for the structure conformation with the minimal potential. ProFOLD reports the structural conformation with sufficiently low potential as the final prediction result (shown in the lower-right corner of Fig. 2), which has a TMscore of 0.84 with the native structure.

The core of our ProFOLD approach is CopulaNet, a deep neural network specially designed to learn inter-residue distances directly from MSA. CopulaNet achieve this objective using three key modules, namely, *MSA encoder, coevolution aggregator*, and *distance estimator*, which are described as below.

*MSA encoder* aims to model the mutations of each residue in the target protein. Here, we represent the MSA with *K* homologue proteins as *K* alignments, each of which consists of a homologue protein aligned with the target protein. Based on each individual alignment, the MSA encoder identifies the mutations of each residue of the target protein, and embeds the mutations into a vector of 64 features.

As a residue’s mutation is highly related to its neighboring residues, the encoder needs to consider a residue of interest together with its neighbors. For this end, we design the encoder as 1D convolutional residual network [18], which uses multiple convolution layers to embed a residue together with its neighbors.

*Coevolution aggregator* aims to measure the residue comutations. For any two residues, the aggregator first calculates the outer product of their embedding features. As the embedding features of a residue encode its mutations, the outer product of two residues’ embedding features could effectively measure the strength of co-mutations between them. Next, to calculate the average outer product obtained from all homologue proteins, we introduce an average-pooling layer into the aggregator. Further details of outer product and average-pooling are shown in the Methods section and Supplementary Figure 2.

*Distance estimator* aims to estimate inter-residue distances according to the acquired residue co-evolutions. Previous studies have revealed structure-related patterns existing in the inter-residue distances. Specifically, two contacting parallel *β*-strands often form a diagonal line, whereas two contacting anti-parallel *β*-strands form an anti-diagonal line. In contrast, two contacting helices usually form a dashed line [19]. Here, we apply a 2D-ResNet to learn these patterns, and thereafter assign these patterns to the estimated inter-residue distances.

As performed by trRosetta [7], we divide the distance range into 37 intervals, i.e., (0Å, 2.5Å), (2.5Å, 3.0Å), (3.0Å, 3.5Å), …, (19.5Å, 20.0Å), and (20.0Å, +∞). This way, we transform the distance estimation problem into a classification problem, which predicts distance interval for each two residues. As results, we obtain a distance distribution over 37 intervals instead of a single estimated distance value (see Supplementary Figure 1 for examples).

### Estimating inter-residue distances using CopulaNet

Using CopulaNet, ProFOLD estimated inter-residue distances for all 104 CASP13 protein domains. For the sake of fair comparison, we evaluated these estimations in terms of precision of the predicted inter-residue contact rather than inter-residue distances. Specifically, for two residues, we summed up the predicted probabilities for their distance below 8 Å, and used the sum as the predicted probability for the two residue being in contact. As shown in Figure 3, on the 31 FM domains, ProFOLD achieved prediction precision of 0.808, 0.673 and 0.536 for top *L*/5, *L*/2 and *L* long-range residue contacts, which is significantly higher than A7D (AlphaFold), the winner group of CASP13, by 0.117, 0.100 and 0.088, respectively. On the other 61 TBM and 12 FM/TBM domains, ProFOLD also outperformed the state-of-the-art approaches (see further details in Supplementary Table 1).

**Figure 3.**
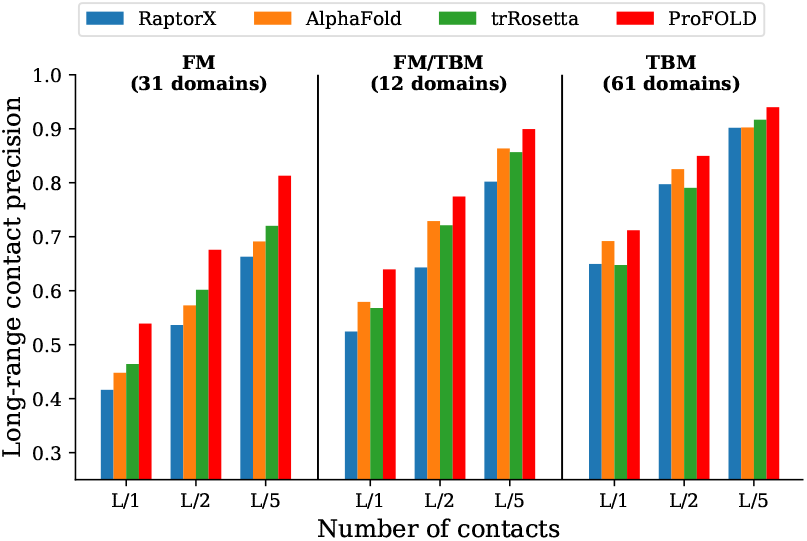
Precision of the predicted inter-residue contacts. Here, we show the precision of the top *L*/5, *L*/2 and *L* long-range residue contacts, where *L* represents protein length. Here the phrase “long-range” refers to two residues with sequence separation over 24 residues. For all CASP13 target proteins, ProFOLD outperformed the state-of-the-art approaches. In particuler, for the 31 FM domains, ProFOLD achieved precision of 0.808, 0.673 and 0.536 for top *L*/5, *L*/2 and *L* long-range residue contacts, which is significantly higher than AlphaFold, by 0.117, 0.100 and 0.088, respectively

We further analyzed the contributions of CopulaNet’s components for estimating inter-residue distances. As mentioned above, the uniqueness of CopulaNet lies at the use of a learnable “encoder and aggregator” framework, rather than traditional statistical models, to model residue co-evolutions. The obtained residue co-evolutions are further fed into a 2D ResNet to assign the structure-related patterns to inter-residue distances. To examine the contributions by these components, we disabled the 2D ResNet in ProFOLD and thus obtained a variant called *ProFOLD w/o R*.

As shown in Figure 4, even using the “encoder and aggregator” framework alone, the variant *ProFOLD w/o R* still showed a high precision (0.367) of top *L* predicted inter-residue contacts on the CASP13 FM targets. The application of the 2D ResNet in ProFOLD further improved the precision by 0.169. These results clearly suggested that the main contribution to the estimation of inter-residue distances comes from the learnable “encoder and aggregator” framework.

**Figure 4.**
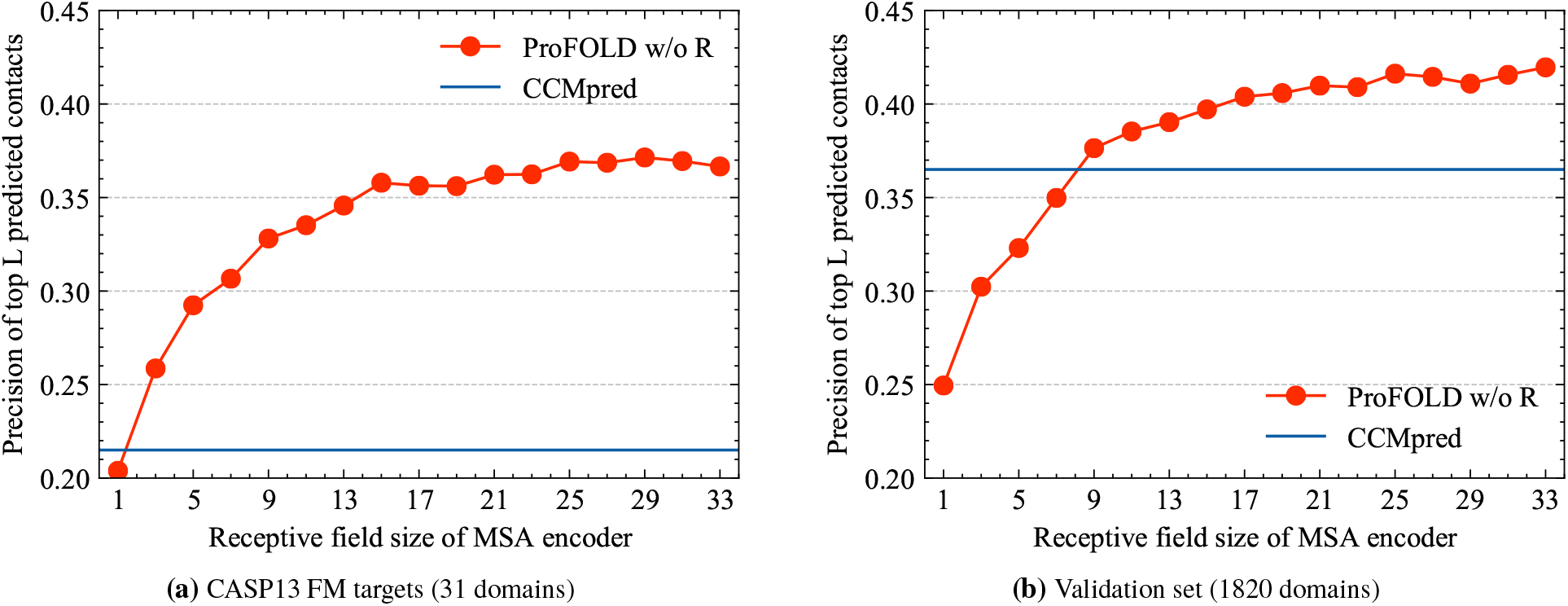
Precision of the predicted inter-residue contacts by the variant *ProFOLD w/o R*. (a) For the 31 CASP13 FM targets, the precision increases with the receptive field size and finally reaches 0.367. (b) On the validation set with 1820 proteins, the precision also increases with the receptive field size and finally reaches 0.420. Even using the “encoder and aggregator” framework alone, the variant *ProFOLD w/o R* still outperformed CCMpred on the two datasets (0.215 and 0.365, respectively)

In addition, on both CASP13 FM targets and validation dataset, the performance of *ProFOLD w/o R* increases with the receptive field size, implying that encoding more neighbors surrounding residues will greatly facilitate distance estimating. In the study, we used an MSA encoder with a receptive field size of 33 (16 1D convolution layers with kernel size 3).

### Predicting protein tertiary structures using ProFOLD

We applied ProFOLD to predict protein tertiary structures and compared it with the state-of-the-art approaches including AlphaFold (A7D group in CASP13) [6], trRosetta[7], top server groups, and top human groups reported by the CASP13 organizer. The prediction results of AlphaFold, top human groups and top server groups were downloaded from CASP13 data archive (https://predictioncenter.org/download_area/CASP13/predictions_trimmed_to_domains/). The prediction results of trRosetta were obtained through re-running it using identical MSA to ProFOLD. The details of these prediction results are summarized in Supplementary Figure 3.

As shown in Figure 5a and Supplementary Table 2, on the 31 FM CASP13 datasets, ProFOLD outperformed the state-of-the-art approaches. Specifically, when setting the cut-off of high-quality structures as TMscore over 0.70, ProFOLD predicted high-quality structures for 18 out of the 31 domains, whereas AlphaFold and trRosetta predicted high-quality structure for only 12 and 7 domains, respectively. Moreover, the average TMscore of ProFOLD’s prediction results is 0.658, which is much higher than that of trRosetta (0.582) and A7D (0.580). Head-to-head comparison clearly demonstrated the advantages of ProFOLD over AlphaFold: for 24 out of the 31 FM domains, ProFOLD outperformed AlphaFold (Figure 5b). ProFOLD also outperformed trRosetta on these FM targets (Supplementary Fig. 4).

**Figure 5.**
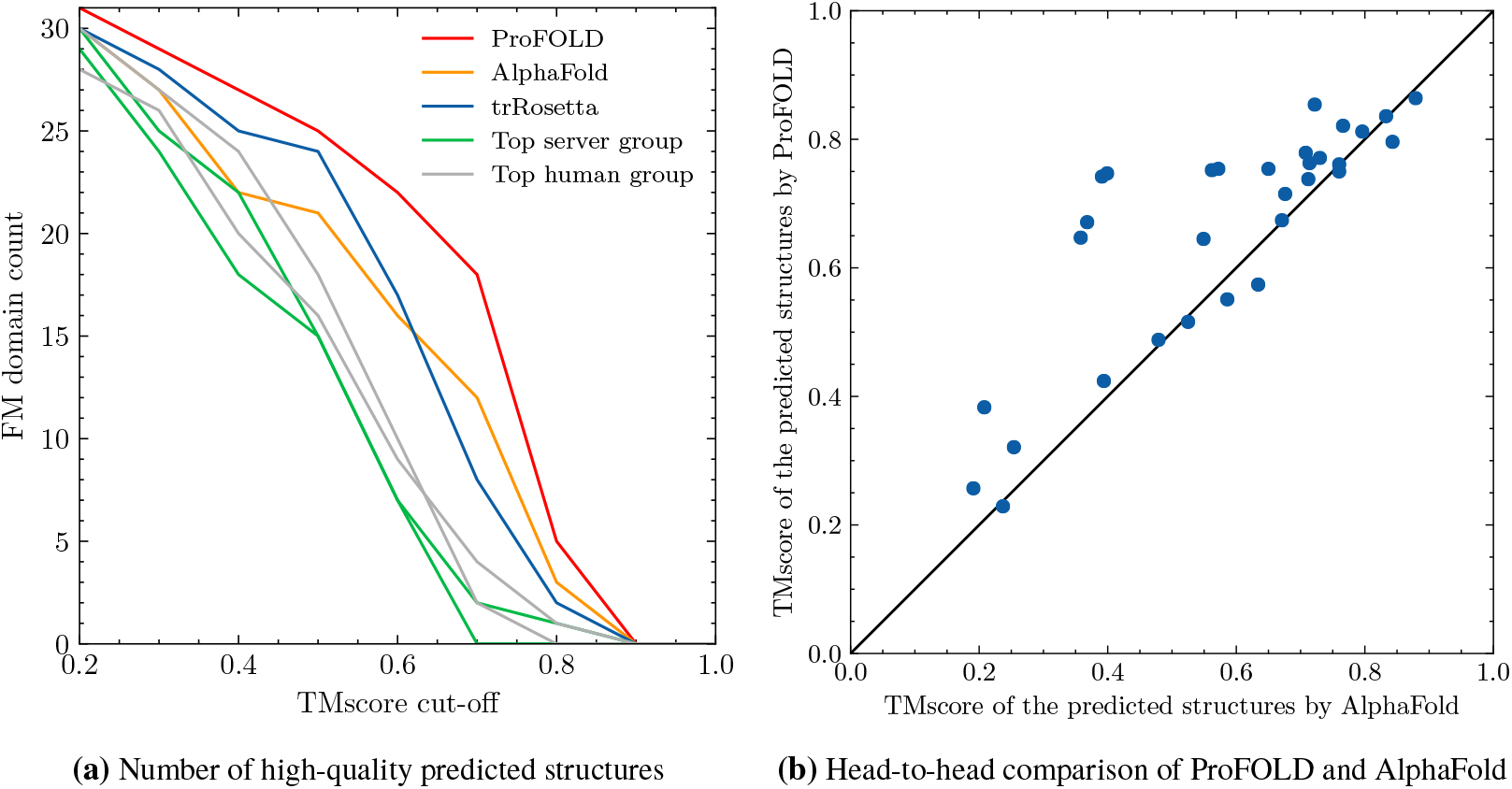
Quality of the predicted tertiary structures for CASP13 FM target proteins. (a) ProFOLD predicted more high-quality structures than the state-of-the-art approaches. When setting the cut-off of high-quality structures as TMscore over 0.70, ProFOLD predicted high-quality structures for 18 out of the 31 domains, whereas AlphaFold and trRosetta predicted high-quality structure for only 12 and 7 domains, respectively. (b) Head-to-head comparison clearly demonstrated the advantages of ProFOLD over AlphaFold: for 24 out of the 31 FM domains, ProFOLD outperformed AlphaFold

We also evaluated ProFOLD on the 61 TBM and 12 TBM/FM target proteins. For these proteins, although similar template structures are available, ProFOLD predicted their structures in pure *ab initio* mode without any reference to the template structure information. As shown in Supplementary Table 2, for these targets, the average TMscore of ProFOLD’s prediction results is 0.785, which is extremely close to the state-of-the-art template-modeling approach Zhang-server (0.787). These results clearly illustrated that the structural information carried by templates might not be necessary for protein structure prediction. Using the accurate estimation of inter-residue distances by CopulaNet, ProFOLD could predict high-quality protein structures without aids of template structures.

We further examined the possible factors that may affect the successful application of ProFOLD. Previous studies have already shown that the quality of predicted structures is highly related to *M*_*eff*_, the number of effective homologue proteins recorded in MSA. As shown in Figure 6a, the correlation coefficient between *M*_*eff*_ and the quality of predicted structures by ProFOLD is as high as 0.69. Therefore, as long as the *M*_*eff*_ of a target protein exceeds 20, TMscore of the predicted structure for this protein is expected to be over 0.60 with high confidence. For proteins T0953s2-D3, T0981-D2, T0991-D1, and T0998-D1, ProFOLD could not predict high-quality structures. The reason might be the fact that for these proteins, *M*_*eff*_ is as small as less than 20. How to improve CopulaNet and ProFOLD to suit the MSAs with only a few homologue proteins remains a future study.

**Figure 6.**
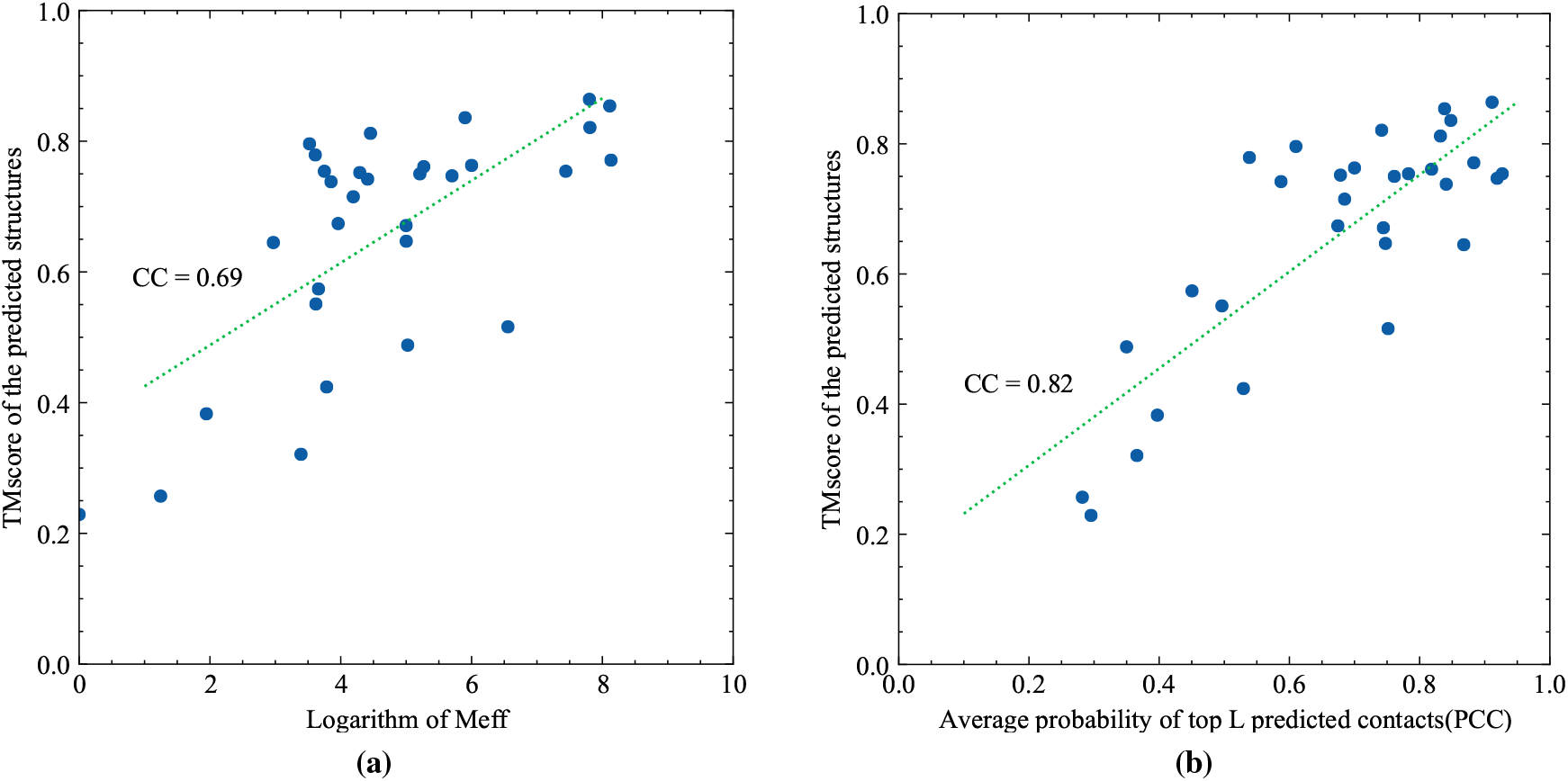
Correlation between quality of the predicted structures and (a) *M*_*eff*_, (b) the average probability of top *L* predicted contacts (PPC). For the CASP13 FM target proteins, the correlation coefficient between *M*_*eff*_ and TMscore of the predicted structures by ProFOLD is as high as 0.69. The correlation efficient between PPC and TMscore of the predicted structures is 0.82.

When the native structure of a target protein is already known, we can easily measure a predicted structure through comparing it with the native structure. However, thing will become challenging when the native structure is not available. For ProFOLD, an interesting and important question is whether we can judge the quality of its prediction results in advance. Here, we provide an effective measure, the average probability of top *L* predicted contacts (denoted as PPC), which is calculated from the estimated inter-residue distances. As shown in Figure 6b, the correlation efficient between PPC and the TMscore of the predicted structures is 0.82. This strong correlation enables us to judge the quality of predicted structure in advance. Specifically, for the target proteins with PPC over 0.60, the TMscore of their predicted structures are expected to be over 0.60 with high confidence.

### Contribution analysis of ProFOLD’s components

To better understand the contribution of the ProFOLD components, we built variants of ProFOLD through disabling each component individually and then compared these variants with ProFOLD. In particular, we first disabled the 2D ResNet in distance estimator and thus obtained a variant called *ProFOLD w/o R*. Next, we further disabled the MSA encoder and obtained another variant called *ProFOLD w/o E+R*. Without MSA encoder to consider the neighboring residues surrounding a residue, *ProFOLD w/o E+R* captures the correlation between two residues without consideration of other residues; thus, *ProFOLD w/o E+R* has roughly the same power to the covariance matrix for distance estimation.

Using protein T1022s1-D1 as an example, we showed the qualitative comparison of the variants in Figure 7. When neither MSA encoder and 2D ResNet is used, the variant *ProFOLD w/o E+R* performed poorly and failed to generate high-quality distance estimations. This result is consistent with the previous observation on the low performance of the approaches based on covariance matrix [11]. When equipped with the MSA encoder module, the variant *ProFOLD w/o R* could generate relatively accurate distance estimations. When both MSA encoder and 2D ResNet are used, ProFOLD gave distance estimations extremely close to the real distance values. These results emphasized the importance of considering neighboring residues in encoding step as well as using the 2D ResNet to learn structure-related patterns existing in the inter-residue distances.

**Figure 7.**
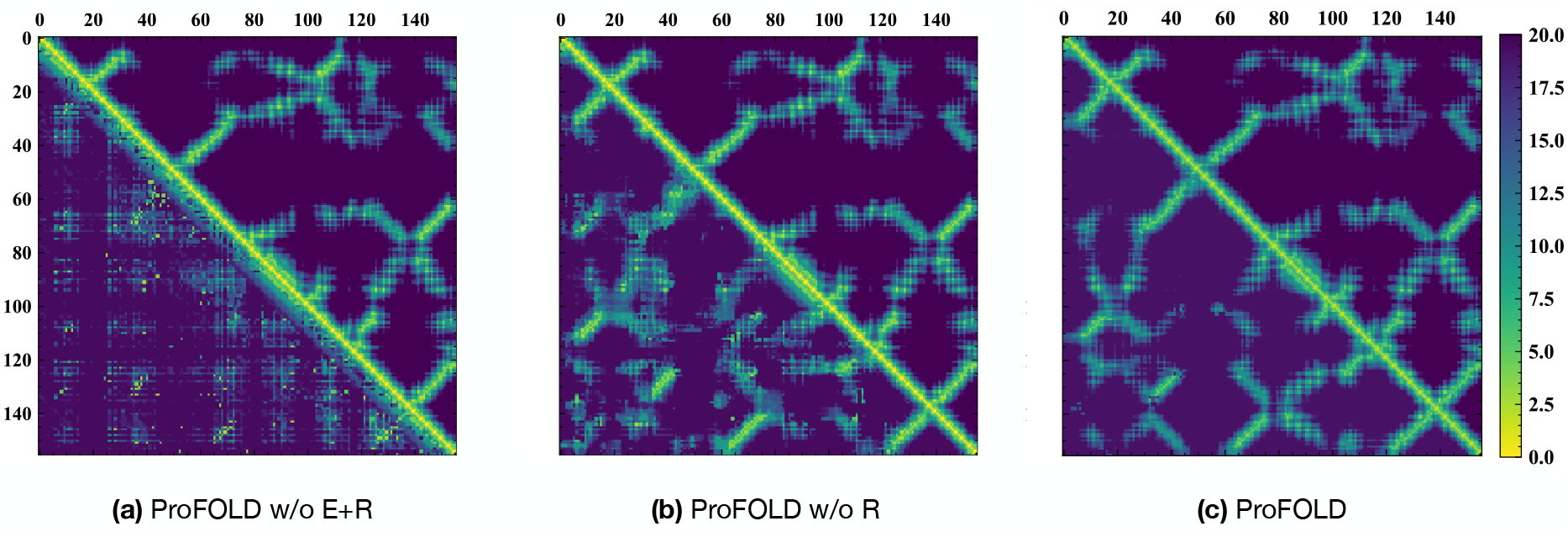
Comparison of the predicted inter-residue distances (bottom left) with the ground-truth distances (upper right) for protein T1022s1-D1. (a) *ProFOLD w/o E+R* performed poorly and failed to generate high-quality distance estimations. (b) When equipped with the MSA encoder module, the variant *ProFOLD w/o R* could generate relatively accurate distance estimations. (c) When both MSA encoder and 2D ResNet are used, ProFOLD gave distance estimations extremely close to the real distance values

To investigate the effect of outer product in coevolution aggregator, we built a variant of ProFOLD (called *ProFOLD w/o OP*) through disabling the outer product operation. Specifically, we modified *h* (*i, j*) in equation (2) to be the concatenation of only *f* (*i*) and *f* (*j*) without *g* (*i, j*). As shown in Supplementary Figure 5, for the short-range residue contacts (between two residues with sequence distance from 6 to 11 residues), *ProFOLD w/o OP* showed roughly the same prediction precision as ProFOLD. This is reasonable as the convolution modules in MSA encoder has already effectively modeled the short range relationship. In contrast, for the long-range residue contacts, the prediction accuracy of *ProFOLD w/o OP* decreased sharply to be significantly lower than ProFOLD. This result clearly demonstrated the importance of the outer product operation in modeling the long-range residue contacts, which cannot be achieved using the convolutional network alone.

### Efficiency of ProFOLD for protein structure prediction

As described above, ProFOLD learns residue co-evolutions directly from MSA rather than the handcrafted features such as covariance matrix. An MSA might have ten of thousands of homologue proteins, whereas the size of covariance matrix is fixed and only determined by the target protein length. Thus, it is interesting to investigate whether ProFOLD could accomplish protein structure prediction within reasonable time on an average computer.

The CopulaNet uses *MSA encoder* and *coevolution aggregator* to process homologue protein sequences. Unlike the final distance estimator processing 2D information of inter-residue coevolution, *MSA encoder* processes 1D sequences only. Moreover, the outer product operations could be efficiently accomplished using the fast matrix multiplication provided by the existing deep neural network frameworks [20]. Thus, compared with distance estimator, both *MSA encoder* and *coevolution aggregator* modules use only a small amount of computations, making the entire running time insensitive to the number of homologue proteins.

Moreover, CopulaNet processes each homologue protein individually; thus, the number of homologue proteins in MSA has little effect on the amount of computer memory required for training and predicting.

As results, for target proteins with less than 500 residues, ProFOLD could finish the whole structure prediction process within 3 hours on an average laptop computer (Intel CPU 2.8G Hz, 16G memory).

## Conclusion

The results presented here for protein structure prediction using the ProFOLD approach have highlighted the special features of learning residue co-evolutions directly from MSA. The abilities of our approach have been clearly demonstrated using CASP13 target proteins as representatives with improved quality of the predicted structures. Using CopulaNet to model the conditional joint-residue distribution, ProFOLD could accurately estimate the inter-residue distances and thereafter predict protein structures. The improved efficiency of ProFOLD is an additional advantage, due mainly to the succinct architecture of CopulaNet. It should also be mentioned that the basic idea and architecture of CopulaNet can be readily modified to calculate conditional joint distribution in other fields besides residue co-evolution.

Although in the proof-of-concept study we demonstrated the application of CopulaNet in *ab initio* prediction of protein structures, the estimated inter-residue distances could also be used to assist template-based prediction approaches. For example, DeepThreader[21] improves threading by incorporating inter-residue distances into scoring function. EigenThreader[22] and CEThreader[23] align target proteins with templates by considering eigenvector decomposition of the predicted inter-residue contacts. These approaches might benefit from the accurate estimation of inter-residue distances provided by CopulaNet.

As CopulaNet attempts to learn residue co-evolution from MSA, it requires that MSA should have sufficient homologue proteins. For the MSAs with only a few homologue proteins, CopulaNet usually cannot accurately estimate inter-residue distances. How to reduce the requirement of the number of homologue proteins remains a future study.

Theoretical analysis suggests a possible failure case of our approach. Consider three residues *r*_*i*_, *r* _*j*_, and *r*_*k*_ in the target protein, where both *r*_*i*_ and *r* _*j*_ are in contact with *r*_*k*_ but there is no contact between *r*_*i*_ and *r* _*j*_. If the sequence distance between *r*_*i*_ and *r*_*k*_ (and between *r* _*j*_ and *r*_*k*_) is sufficiently long, MSA encoder cannot perfectly model the effect of *r*_*k*_ on *r*_*i*_ and *r* _*j*_, thus perhaps causing ProFOLD to incorrectly report a contact for residue *r*_*i*_ and *r* _*j*_. The increase of receptive field size in MSA encoder will partially alleviate this problem; however, when receptive field size is already large, further increase of it will bring limited gains. A perfect way to model long-distance influence among residues is another future study.

In summary, our work on learning residue co-evolution directly from MSA together with recent developments in constructing high-quality MSAs will undoubtedly contributed to more accurate prediction of protein tertiary structures.

## Methods

### Architecture of CopulaNet

CopulaNet consists of three key modules, i.e., *MSA encoder, co-evolution aggregator*, and *distance estimator*.

*MSA encoder* embeds residue mutations using a 1D convolutional residual network [18]. The residual network has 8 residual blocks, and each residual block consists of two batch-norm layers, two 1D convolution layers with 64 filters (with kernel size of 3) and exponential linear unit (ELU) [24] non-linearities (Supplementary Figure 6).

*Co-evolution aggregator* measures the co-mutations between two residues. Before presenting the design of *co-evolution aggregator* module, we describe the notations first.

Consider a target protein with *L* residues *t*_1_*t*_2_ … *t*_*L*_, and a pre-built MSA containing *K* homologue proteins. By applying *MSA encoder* on the *k*-th homologue protein in MSA, we obtain a total of *C* × *L* embedding features, denoted as *X*_*k*_ ∈ ℝ^*C*×*L*^, where *C* represents the number of output channels of *MSA encoder*. For residue *t*_*i*_ in the target protein, its embedding features calculated from all homologue proteins are aggregated together. The aggregated embedding features, denoted as *f* ∈ ℝ^*C*×*L*^, are calculated as follows.

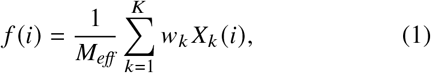

where *w*_*k*_ denotes the weight of the *k*-th homologue protein, and 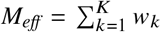 represents the sum weight of all homologue proteins. Following the convention [14], we calculate the weight *w*_*k*_ as the inverse of the number of similar homologue proteins that share at least 80% sequence identity with the *k*-th homologue, and thus *M*_*eff*_ represents the number of effective homologue proteins in the MSA.

For two residues *t*_*i*_ and *t* _*j*_ in target protein, the co-evolution aggregator measures their co-mutations using aggregated co-evolution features *h* (*i, j*) ∈ ℝ^*D*^, where *D* denotes the number of output channels of *co-evolution aggregator. h* (*i, j*) is a concatenation of the aggregated embedding features and their outer products:

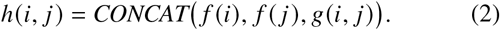

Here, *g* (*i, j*) ∈ ℝ^*C*×*C*^ represents the aggregated outer products of the embedding features for residue *t*_*i*_ and *t* _*j*_, which is calculated as below.

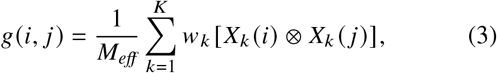

where “⊗” represents the outer product operation.

To summarize, the aggregated co-evolution features consists of *C* × 2 aggregated embedding features and *C* × *C* aggregated outer product features. In this study, the output channel size *C* of *MSA encoder* is set as 64. Thus, the co-evolution aggregator generates a total of 4224 (64 × 2 + 64 × 64) aggregated co-evolution features for any two residues in target protein. An example of the outer product operation is shown in Supplementary Figure 2.

*Distance estimator* aims to estimate inter-residue distances according to the obtained residue co-evolutions using a 2D-ResNet with 72 residual blocks. Each block consists of two batch-norm layers, two 2D dilated convolution layers, and exponential linear unit (ELU) nonlinearities.

The further details of CopulaNet, including hyperparameters, and training process, are provided in Supplementary text.

### Approximating the conditional joint-residue distribution using CopulaNet

Consider a target protein with *L* residues. The alignment of a homologue protein with target protein might have insertions, deletions, and mutations. As our objective is to describe residue co-mutations, we follow the conventions to discard the insertions [9]. Thus, a homologue protein can be represented as a *L*-long sequence composing of residues and a special character ‘-’ representing deletion. Assume each homologue protein is a random variable = *r*_1_*r*_2_ … *r*_*L*_ sampled from the joint distribution *P* (*r*_1_, *r*_2_, …, *r*_*L*_). In this subsection, we will show that for residues *r*_*i*_ and *r* _*j*_, CopulaNet could perfectly approximate the conditional joint-residue distribution *P*(*r*_*i*_, *r* _*j*_ |other residues).

As mentioned above, from the *k*-th homologue protein, the MSA encoder extracts the mutations for the *i*-th residue in the target protein and embeds these mutations into a vector *X*_*k*_ (*i*) using a CNN. Previous studies have already suggested that, although in principle CNN has the ability to describe long range relationship, it focuses on neighboring residues rather than distant pairs [25]. Thus, to be more precise, we rewrite *X*_*k*_ (*i*) as *X*_*k*_ (*i, N*_*i*_), where *N*_*i*_ represents the neighboring residues of the *i*-th residue with significant effects on its mutation. Accordingly, we rewrite the aggregated embedding feature *f* (*i*) as *f* (*i, N*_*i*_), and rewrite the aggregated outer product *g*(*i, j*) as *g*(*i, N*_*i*_, *j, N*_*j*_):

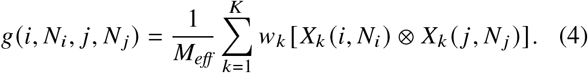

Outer product could effectively capture the correlation between two variables. To understand how outer product works, we showed in Supplementary Figure 2 the outer product of two one-hot feature vectors as an illustrative example. In this case, the 3rd entry of the one-hot vector *f* (*i, N*_*i*_) is 1, and the 1st entry of the one-hot vector *f* (*j, N*_*j*_) is 1. Thus, the outer product of them has 1 at the (3, 1) -th entry, which clearly reveal the correlation between these vectors. In practice, instead of the simple one-hot feature vectors, more informative real-value feature vectors are used.

Through averaging over all homologue proteins, the aggregated outer product *g* (*i, N*_*i*_, *j, N*_*j*_) could perfectly approximate the joint distribution *P* (*x*_*i*_, *N*_*i*_, *x*_*j*_, *N*_*j*_). Similarly, CopulaNet also has the ability to approximate the joint distribution *P* (*N*_*i*_, *N*_*j*_). The ratio of these two joint distributions provides a perfect approximation to the conditional joint-residue distribution *P* (*x*_*i*_, *x*_*j*_ |*N*_*i*_, *N*_*j*_). The approximation extent will be increased when using large neighborhood (See Results and discussion section for details).

### Benchmark dataset

In this study, we prepared a benchmark dataset (containing 31,247 protein domains) using the same pipeline to AlphaFold [6]. We randomly divided this dataset into two parts, a training dataset containing 29,427 proteins, and a validation dataset containing 1,820 proteins.

We tested our methods on CASP13 targets, which consists of 104 domains derived from 71 official targets (the first target was released on May 1, 2018). The 104 domains are officially split into three categories: FM (31 domains), FM/TBM (12 domains) and TBM (61 domains).

### MSA generation and representation

ProFOLD takes multiple sequence alignment as its only input. For a target protein, we first search its homologue proteins by running DeepMSA [26] (with default parameters) against sequence databases uniclust30 (version 2017-10), uniref90 (version 2018-03) and metaclust50 (version 2018-01). All these sequence databases were released before independent test sets and thus there is no overlap between them with the test dataset.

In the study, we represent the obtained MSA as a collection of sequence pairs. Each sequence pair contains the target protein and a homologue protein. We construct two equal-length strings by adding gaps in aligned sequences so that matching characters are aligned in successive positions (Fig. 2). Then we encode each position with a binary vector of 41 elements, including 20 elements for target protein and 21 elements for homologue protein. Here, 20 elements are one-hot vector that represents 20 amino acid types, and a special character ‘-’ is introduced to represent gaps.

### Structure determination using distance potential

In the study, we build protein tertiary structures from the predicted inter-residue distances in a similar way to trRosetta[7]. Specifically, we first convert the estimated inter-residue distance distributions into a smooth potential function using the DFIRE [27] paradigm. Then, we use *MinMover* in PyRosetta[28] to search for the tertiary structure with the minimal potential, yielding (centroid) models. Finally, these coarse-grained models are refined into full-atom models by executing *FastRelax* in Rosetta.

## Data availability

The datasets used in this study is available via http://protein.ict.ac.cn/ProFOLD.

## Code availability

We developed a web server that is available through http://protein.ict.ac.cn/ProFOLD. All source codes and models of ProFOLD are publicly available through http://protein.ict.ac.cn/ProFOLD.

## Acknowledgements

We would like to thank the National Key Research and Development Program of China (2018YFC0910405), and the National Natural Science Foundation of China (31671369, 31770775) for providing financial supports for this study and publication charges.

## Author contributions

FJ, JZ and DB conceived the study. FJ designed and implemented the neural network, and performed the computation. FJ, JZ, BS, TL, WZ, and DB analyzed the experimental results. FJ, LK and DB established the mathematical framework. FJ and DB wrote the manuscript. All authors read and approved the final manuscript.

## Additional information

Supplementary information is available for this paper at https://.

Correspondence and requests for materials should be addressed to.

Reprints and permissions information is available at http://www.nature.com/reprints.

