## Supplemental tables and figures for "CopulaNet: Learning residue co-evolution directly from multiple sequence alignment for protein structure prediction"

6 October 2020

### Supplementary Material

**Supplementary Text**

**Supplementary Table 1:** Precision of the predicted long-range contacts over CASP13 targets

**Supplementary Table 2:** Quality of the predicted tertiary structures for CASP13 targets. Here, we list TMscore of top1/top5 predicted structure

**Supplementary Figure 1:** Predict tertiary structure for CASP FM target protein T0992-D1 using ProFOLD

**Supplementary Figure 2:** Outer product of the embedding features and average pooling in the coevolution aggregator module

**Supplementary Figure 3:** Quality of the predicted structure for the CASP13 FM targets

**Supplementary Figure 4:** Head-to-head comparison of trRosetta and ProFOLD on the CASP13 FM targets

**Supplementary Figure 5:** Precision of the predicted inter-residue contacts by ProFOLD and the variant *ProFOLD w/o OP*
**Supplementary Figure 6:** The components of residual block used in *MSA encoder* and *distance estimator*

### Supplementary Text

#### Hyperparameters of the neural networks used by CopulaNet

- 1D ResNet: 8 one-dimensional blocks with 64 channels.
- 2D ResNet: 18 groups of 4 two-dimensional blocks with 96 channels, cycling through dilations 1, 2, 4, 8.
- Optimizer: Adam with learning rate $1e^{-3}$.
- Batch: mini-batch of 3 crops on each of 4 GPU workers.
- Nonlinearity: ELU.
- Loss: cross-entropy.
- Training time: about 15 hours for 60,000 steps.

#### Evaluation criteria of the predicted structures and inter-residue distances

We evaluate the quality of the predicted structures by aligning them with the native structure using DeepAlign [1]. For the CASP13 targets, DeepAlign reports TMscore [2] of the predicted structures, which are close to the TMscore published by CASP13 organizer. For each target protein, we predict multiple tertiary structures (called models). Following the CASP convention, we evaluate the prediction approaches in terms of both top 1 and top 5 models.

For the sake of fair comparison, we evaluate the estimated inter-residue distances through transforming into inter-residue contacts. Specifically, for two residues, we sum up the predicted probabilities for their distance below 8Å, and use the sum as the predicted probability for the two residue being in contact. We evaluate the precision of the top $L/5$, $L/2$ and $L$ predicted contacts, where $L$ denots the length of protein sequence [3]. We focus on the long-range contacts (between two residues with sequence separation ≥24 residues), because they are harder to predict and more important for constructing structure.

### Supplementary tables

**Table 1 |** Precision of the predicted long-range contacts over CASP13 targets.

| Dataset | FM (31 domains) | | | FM/TBM(12 domains) | | | TBM (61 domains) | | |
|---|---|---|---|---|---|---|---|---|---|
| | L | L/2 | L/5 | L | L/2 | L/5 | L | L/2 | L/5 |
| RaptorX | 0.416 | 0.536 | 0.662 | 0.524 | 0.643 | 0.802 | 0.650 | 0.797 | 0.902 |
| A7D | 0.448 | 0.573 | 0.691 | 0.579 | 0.729 | 0.864 | 0.692 | 0.825 | 0.902 |
| trRosetta | 0.466 | 0.602 | 0.720 | 0.568 | 0.721 | 0.857 | 0.648 | 0.791 | 0.917 |
| ProFOLD | **0.536** | **0.673** | **0.808** | **0.641** | **0.774** | **0.899** | **0.712** | **0.850** | **0.940** |

**Table 2 |** Quality of the predicted tertiary structures for CASP13 targets. Here, we list TMscore of top1/top5 predicted structure

| Methods | All (104) | TBM (61) | FM/TBM(12) | FM (31) |
|---|---|---|---|---|
| A7D | 0.699/0.733 | 0.761/0.786 | 0.691/0.739 | 0.580/0.626 |
| Zhang (Human) | 0.692/0.719 | **0.801**/0.816 | 0.605/0.665 | 0.509/0.549 |
| MULTICOM (Human) | 0.688/0.722 | 0.794/**0.817** | 0.645/0.675 | 0.495/0.551 |
| QUARK | 0.672/0.699 | 0.786/0.808 | 0.589/0.648 | 0.479/0.503 |
| Zhang-Server | 0.671/0.699 | 0.787/0.807 | 0.593/0.627 | 0.475/0.514 |
| RaptorX-DeepModeller | 0.653/0.674 | 0.774/0.786 | 0.561/0.592 | 0.451/0.486 |
| trRosetta | 0.667/0.675 | 0.719/0.727 | 0.622/0.625 | 0.582/0.594 |
| ProFOLD | **0.742/0.748** | 0.785/0.791 | **0.738/0.741** | **0.658/0.666** |

### Supplementary figures

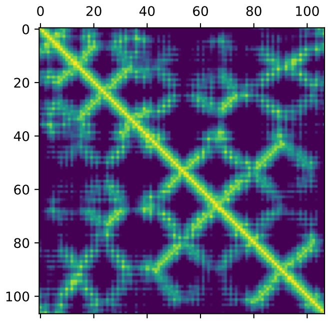


**(a)** Native distance

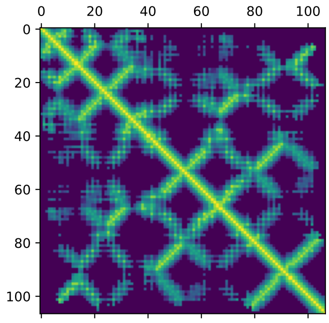


**(b)** Predicted distance

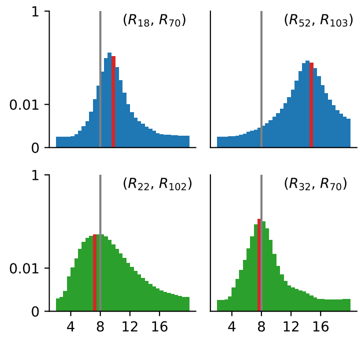


**(c)** Distribution

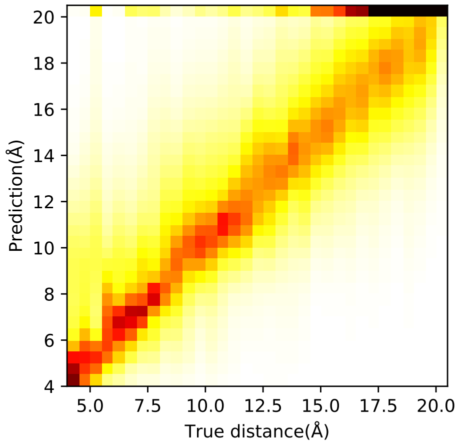


**(d)** Predicted distribution

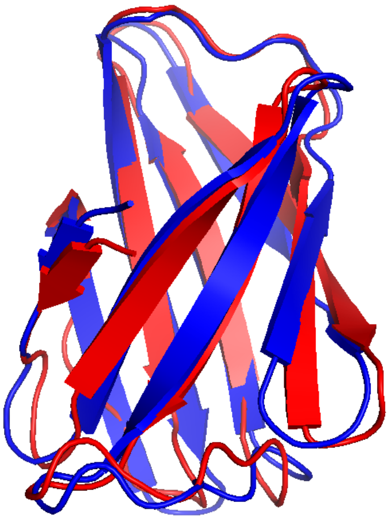

**(e)** Native structure

**Figure 1 | Predict tertiary structure for CASP FM target protein T0992-D1 using ProFOLD.** (a) The native inter-residue distances. (b) The predicted inter-residue distances using CopulaNe, which is very close to the native distances. (c) The predicted inter-residue distance distribution for 4 residue pairs. Here, the bin covering the ground-truth distance is highlighted in red. True-contact distributions are plotted in green, otherwise in blue. The gray line represents the contact cutoff 8Å. (d) The comparison of the predicted distance distributions and the ground-truth distances. The concentration on the diagonal line suggests that the predicted distances are very close to the ground-truth distances. (e) The predicted tertiary structure (in blue) and the native structure (in red). The TMscore between them is 0.84, indicating that the predicted structure has high quality

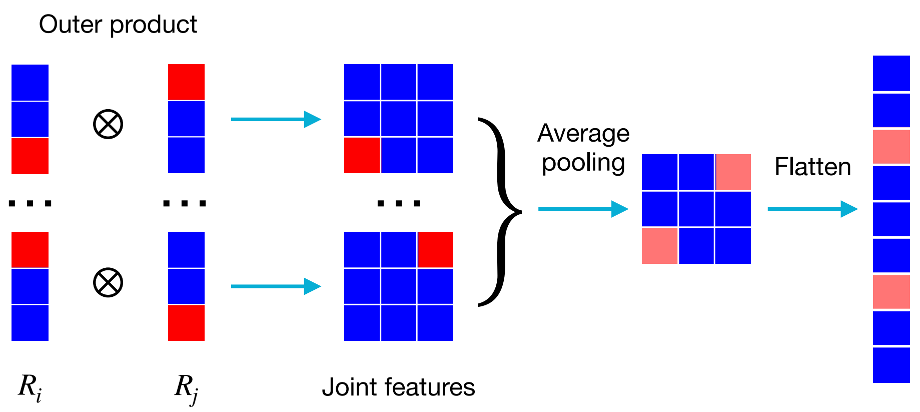


**Figure 2 | Outer product of the embedding features and average pooling in the coevolution aggregator module.** To understand how outer product works, we showed here the outer product of two one-hot feature vectors as an illustrative example. For the 1st homologue protein, the 3rd entry of the one-hot vector for $R_i$ is 1, and the 1st entry of the one-hot vector for $R_j$ is 1. Thus, the outer product of them has 1 at the $(3, 1)$-th entry, which clearly reveal the correlation between these vectors. Next, we use average pooling to combine the outer products calculated for all homologue proteins. In practice, instead of the simple one-hot feature vectors, more informative real-value feature vectors are used.

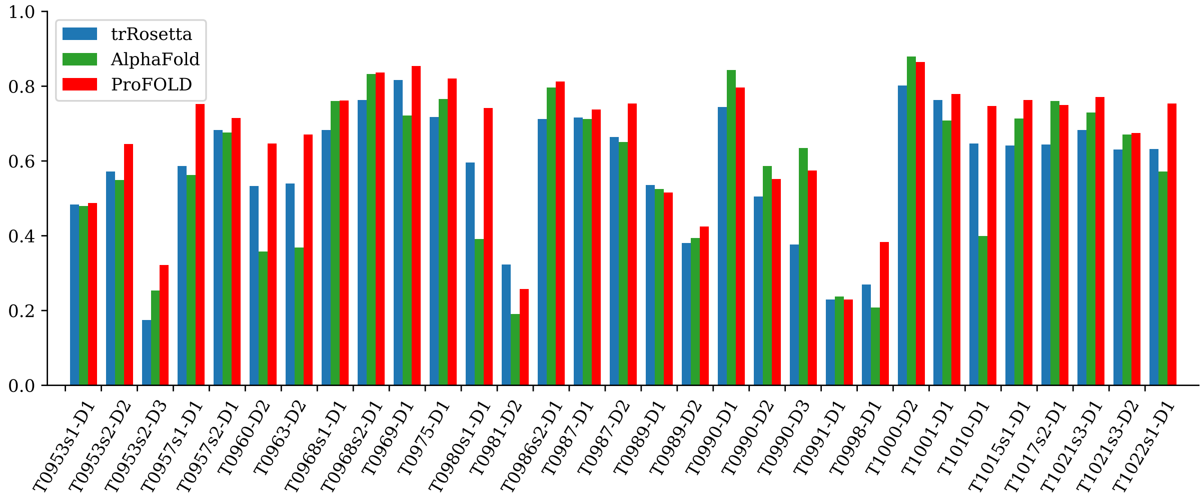


**Figure 3 | Quality of the predicted structure for the CASP13 FM targets.** Here we show TMscore of the top 1 model predicted by trRosetta, AlphaFold and ProFOLD

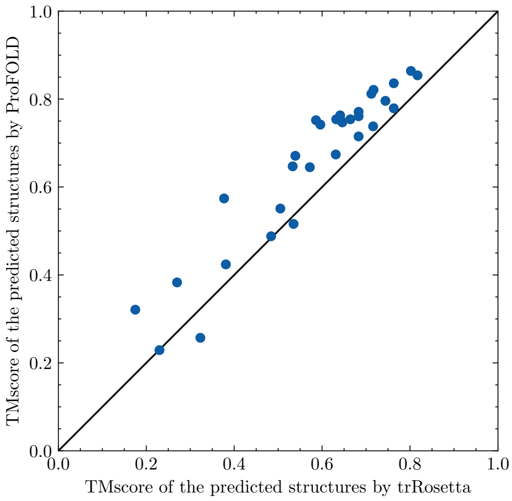


**Figure 4 | Head-to-head comparison of trRosetta and ProFOLD on the CASP13 FM targets**. Here we use TMscore of the top 1 model predicted by the two approaches. For 27 out of the 31 FM targets, ProFOLD outperformed trRosetta

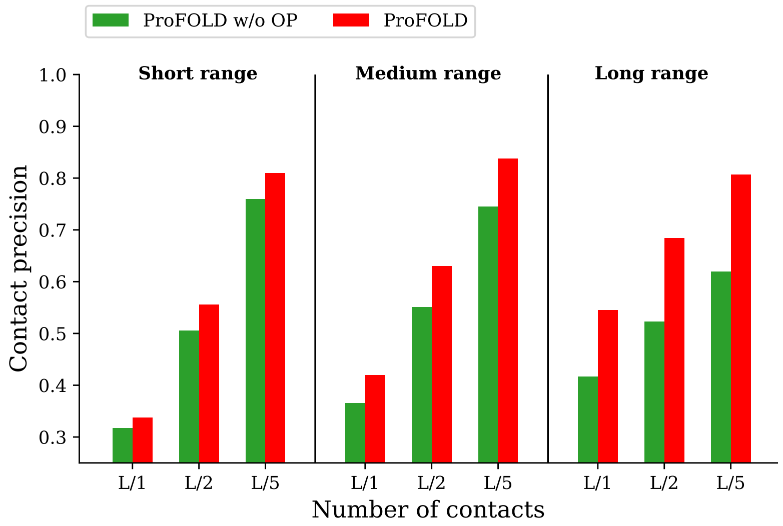


**Figure 5 | Precision of the predicted inter-residue contacts by ProFOLD and the variant** *ProFOLD w/o OP* **on CASP13 targets.** Here, the variant *ProFOLD w/o OP* was built through disabling the outer product operation of ProFOLD. For the short-range residue contacts (between two residues with sequence distance from 6 to 11 residues), *ProFOLD w/o OP* showed roughly the same prediction precision as ProFOLD. This is reasonable as the convolution modules in MSA encoder has already effectively modeled the short range relationship. In contrast, for the long-range residue contacts, the prediction accuracy of *ProFOLD w/o OP* decreased sharply to be significantly lower than ProFOLD. This result clearly demonstrated the importance of the outer product operation in modeling the long-range residue contacts, which cannot be achieved using the convolutional network alone.

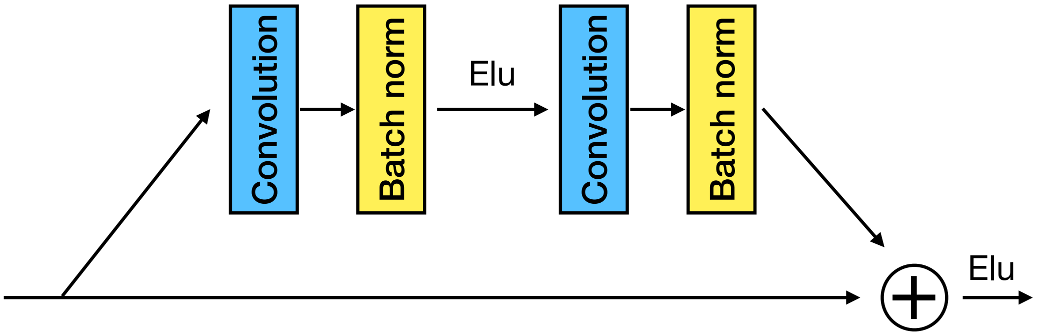


**Figure 6 | The components of residual block used in** *MSA encoder* **and** *distance estimator*. The residual network in *MSA encoder* has 8 residual blocks, and each residual block consists of two batch-norm layers, two 1D convolution layers with 64 filters (with kernel size of 3) and exponential linear unit (ELU) nonlinearities. *Distance estimator* has 72 residual blocks with roughly same components, but use 2D $3 \times 3$ dilated convolution layers with 96 filters.
